# Exogenous auxins for proline regulation in heat-stressed plants

**DOI:** 10.64898/2025.12.20.695708

**Authors:** Abdussabur M. Kaleh, Joann K. Whalen

## Abstract

Microbial indole-3-acetic acid (IAA) has long been recognized as a driver of plant growth and developmental plasticity. Recent studies show that microbial IAA production is sustained or even enhanced at elevated temperatures, suggesting that microbial auxins may contribute to plant thermotolerance by stabilizing auxin signalling and supporting metabolic adaptation. Here, we synthesize emerging molecular and physiological evidence linking microbial IAA to proline turnover during thermomorphogenesis. We propose that microbial IAA establishes a regulatory window in which proline metabolism transitions between early osmoprotective synthesis and later catabolism that fuels elongation and redox balance. This integration involves crosstalk between the HSP90-TIR1 auxin perception module, heat-responsive MPK-IAA8 signalling, mitochondrial redox regulators (SSR1, HSCA2), and hormonal interactions with abscisic acid and ethylene. We outlined four mechanistic hypotheses and associated experiments to test how microbial IAA modulates proline homeostasis. This framework highlights microbial auxins as metabolic integrators during heat stress and provides a conceptual basis for leveraging auxin-producing microbes to enhance plant resilience under global warming.

**Highlight:** Microbial auxins modulate the proline cycle during heat stress by integrating hormonal, redox, and mitochondrial signals, offering a mechanistic framework for how microbial IAA enhances thermomorphogenesis and plant heat resilience.

## Introduction

Auxin regulates almost every aspect of plant growth and development, acting as a link between environmental stimuli and metabolic adaptation. Its role in thermomorphogenesis, the developmental reprogramming that produces elongated petioles, narrower leaves and altered organ orientation under heat stress is well established (Casal and Balasubramanian, 2019). At the molecular level, this process is regulated by the PHYTOCHROME-INTERACTING FACTOR 4 (PIF4) transcription factor (Koini *et al.,* 2009). Under ambient temperatures, phytochrome B remains in its active Pfr form, repressing PIF4; elevated temperatures shift it to the inactive Pr form, relieving this repression. In parallel, heat-induced changes in RNA secondary structure promote translation of PIF7 mRNA, which cooperates with PIF4 to activate auxin biosynthetic genes such as *YUCCA8* and *TAA1* (Kan *et al.,* 2023). The resulting increase in indole-3-acetic acid (IAA) initiates a signaling cascade that drives thermomorphogenic elongation (Gray *et al.,* 1998; Sun *et al.,* 2012).

Auxins also regulate heat tolerance at the cellular level. PIN-LIKES (PILS) proteins, particularly PILS6, act as intracellular auxin carriers that shapes auxin availability and distribution (Suar and Kleine-Vehn, 2019). Heat stress reduces PILS6 abundance, redistributing auxin and increases auxin signaling (Feraru, 2019). Activation of Auxin Response Factors (ARFs) is necessary for these morphological responses, since ARF-deficient plants fail to elongate under elevated temperature (Reed et al., 2018). These processes demand substantial metabolic energy and redox coordination, suggesting that auxin must interface directly with cellular energy pathways. A key node in this connection is Heat Shock Protein 90 (HSP90), which stabilizes the auxin co-receptor TRANSPORT INHIBITOR RESPONSE1 (TIR1) through complex formation with SGT1 (Wang et al., 2016). Upon heat exposure, elevated HSP90 levels enhance TIR1 stability and promote Aux/IAA degradation, facilitating downstream transcriptional responses including *IAA19*, *GH3.17*, and *DR5* activation resulting in a sustained auxin sensitivity even when biosynthesis fluctuates. Interestingly, HSP90-TIR1 acts downstream of auxin biosynthesis, and independently of PIF4-induced *YUCCA8* expression. Inhibition of HSP90 activity (e.g., with geldanamycin) diminishes the expression of auxin-responsive genes even in the presence of IAA. Together, these findings show auxin as the primary hormonal drivers of thermomorphogenesis, with heat-stabilized components like HSP90 ensuring that auxin signaling remains effective and responsive through TIR1 stabilization and enhanced downstream transcriptional activation.

Plants also coexist with IAA-producing microorganisms as endophytes, free-living in the rhizosphere or phyllosphere, including *Bacillus*, *Pseudomonas*, and *Trichoderma* species that continue to synthesize IAA at ≥35 °C (Table 1). Many of these microbes enhance plant heat tolerance, yet their influence on the auxin-driven process of thermomorphogenesis remains largely unexplored. We propose that microbial IAA acts as an auxiliary hormonal input that integrates into host metabolic networks by modulating proline homeostasis, a central hub of osmoprotection, redox buffering, and energy turnover. By shifting the balance between proline synthesis and catabolism, microbial IAA may enable plants to transition from protection to recovery during thermomorphogenesis.

**Table 1.**
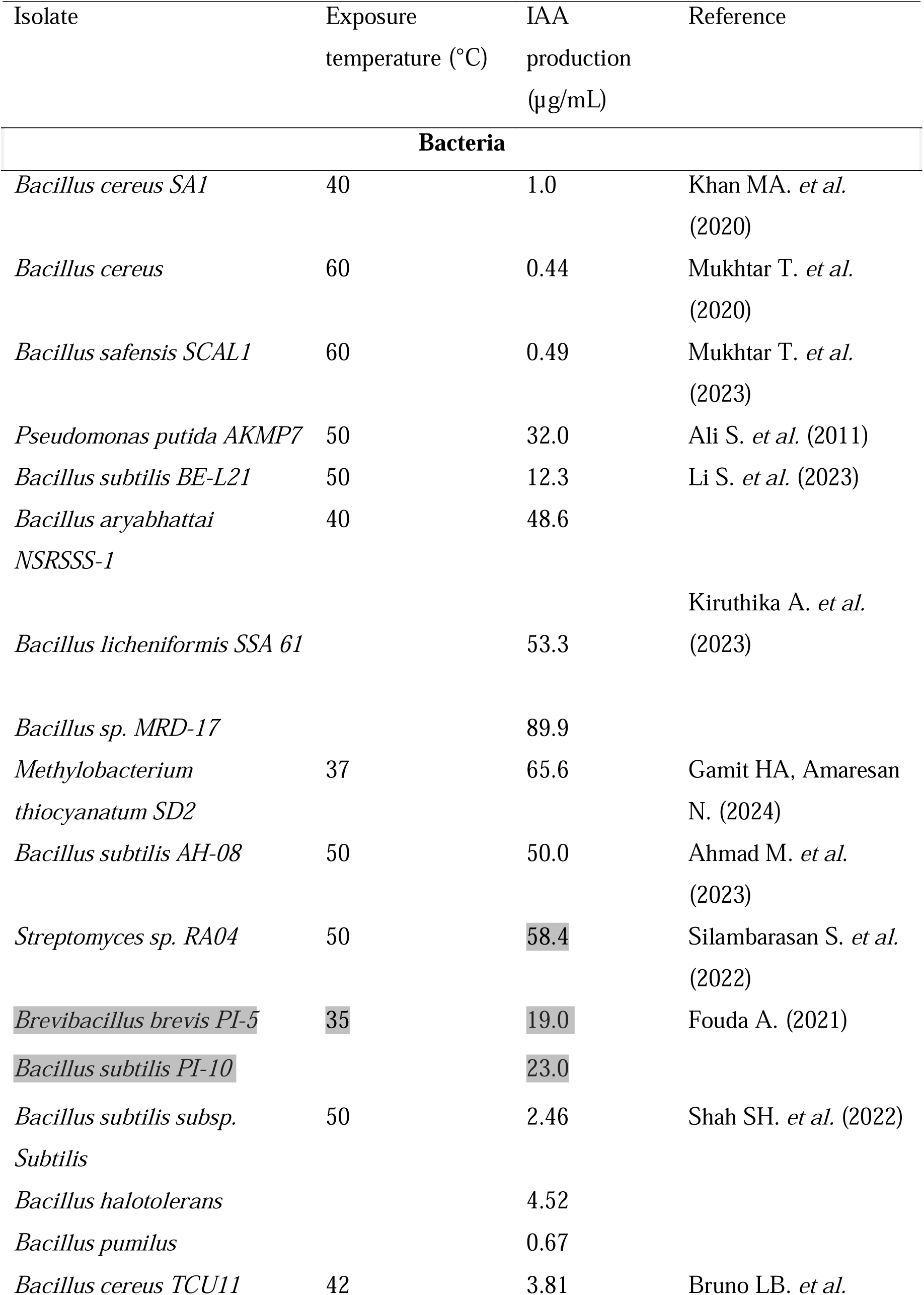
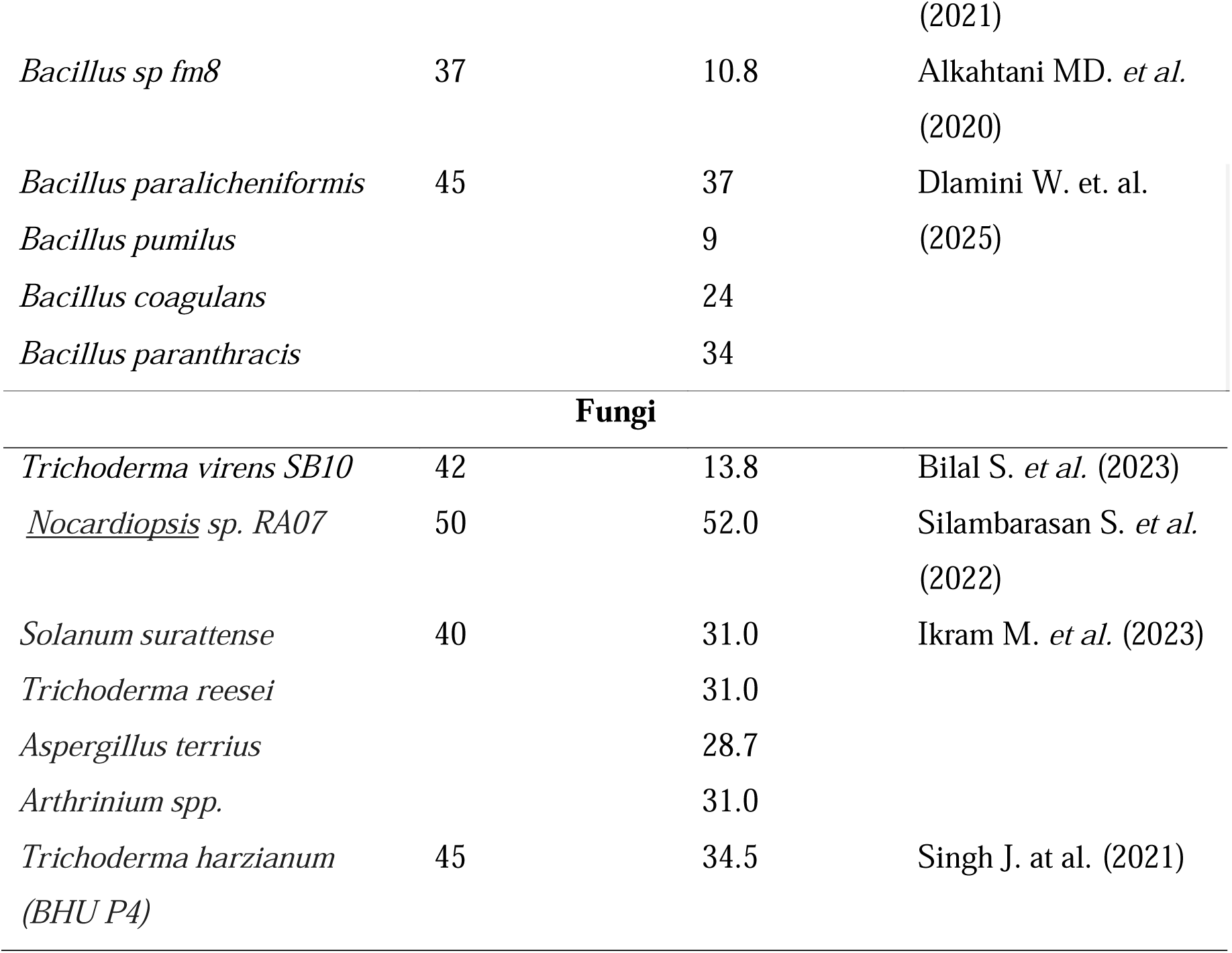
Indole-3-acetic acid (IAA) production by bacteria and fungi at ≥35°C in laboratory cultures.

### How might microbial auxins regulate thermomorphogenic metabolism?

Numerous studies have reported that IAA-producing microbes mitigate heat stress in diverse plant species. Most attribute this resilience to elevated antioxidant activity and proline levels, consistent with auxin-mediated stress protection. However, the reported proline response vary widely between studies: for example, soybean (*Glycine max*) inoculated with IAA-producing microbes under 40°C heat stress for two weeks, showed proline levels ranging from 0.12 to 16.5 μmol g^−1^ FW min^−1^ (Ismail et al., 2018; Ismail et al., 2020), whereas tomato (*Solanuum lycopersicum L.*) cultivars inoculated with IAA-producing microbes under 42°C stress showed values ranging 1.7 to 35.1 μmol g□¹ FW min□¹ (Tripathi et al., 2021; Duc, 2023). Such variability suggests that microbial IAA does not simply elevate proline levels but instead modulates the balance between proline synthesis and degradation depending on physiological context, adjusting metabolic flux to the plant’s developmental state and stress intensity. Thermomorphogenesis involves a metabolic phase shift from early accumulation of osmoprotectants during initial stress to later catabolic energy mobilization that supports renewed cell elongation and growth. Microbial IAA may act as a regulatory signal coordinating this transition. Root-and leaf-associated microbes can produce tens of micrograms of IAA mL□¹ at elevated temperature (Table 1), concentrations sufficient to influence host signaling. And endogenous auxin biosynthesis often declines under heat (Sakata et al., 2010), destabilizing the perception machinery. Therefore, incorporation of microbial IAA into the plant hormone pool could stabilize the HSP90-TIR1 complex, maintaining signal transduction and promoting expression of elongation-associated genes such as *IAA19* and *GH3.17*. In this framework, microbial IAA acts not as an additive hormone but as a stabilizing buffer that preserves developmental-metabolic coherence during heat stress.

Elevated temperature also stimulates microbial IAA biosynthesis, creating positive feedback between microbial metabolism and host demand. Many bacteria and fungi up-regulate IAA synthesis at ≥ 35 °C through both L-tryptophan-dependent and-independent pathways that ensure continuous production even if one route is disrupted (Tang et al., 2023). For example, *Bacillus coagulans* LB6 produced 23 mg IAA mL□¹ after 24 h at 45 °C (Dlamini et al., 2025). Overproduction of IAA in *Sinorhizobium meliloti* 1021 improved survival under heat, osmotic, and UV stress by enhancing biofilm formation and exopolysaccharide synthesis (Imperlini et al., 2009). *Bradyrhizobium japonicum* pre-treated with IAA showed 78 % higher viability after heat shock (Donati et al., 2013), and warming from 30 °C to 37 °C increased IAA output by 50 % in *Azospirillum brasilense* SM and by 18 % in *Bacillus* spp. (Mohite, 2013). Therefore, microbial IAA biosynthesis and microbial thermotolerance are mutually reinforcing, enabling sustained auxin release precisely when host plants require hormonal and metabolic support. Beyond biosynthesis, exogenous IAA also enhances microbial physiological stability. *Azospirillum brasilense* mutants lacking the key biosynthetic enzyme *ipdC* exhibit membrane depolarization and impaired root colonization under heat stress, but supplementation with low doses restores membrane potential and rescues colonization efficiency (Ganusova et al., 2025). These findings indicate that microbial-or plant-derived IAA support both microbial survival and plant auxin signaling, creating a bidirectional benefit. Therefore, microbial auxins mediate a reciprocal adaptation mechanism in which elevated temperature enhances microbial IAA biosynthesis and resilience, while the resulting auxin flux modulates host proline metabolism and developmental plasticity. This mutualistic exchange forms a dynamic feedback system through which plant-microbe partnerships sustain growth under heat stress.

### How could microbial IAA reshape proline homeostasis under heat stress?

Proline metabolism is central for plant thermotolerance. During heat stress, proline contributes to osmotic adjustment, stabilizes proteins and membranes, and scavenges reactive oxygen species (Kavi et al., 2022). Yet excessive proline accumulation can be harmful, Arabidopsis overproducing proline showed reduced survival after heat treatment (Lv et al., 2011). In rice, balanced proline catabolism is equally critical, loss of mitochondrial enzyme *OsProDH* enhanced heat tolerance by preventing ROS over-accumulation, whereas overexpression rendered plants hypersensitive (You et al., 2012). These findings demonstrate that effective thermotolerance depends on a dynamic equilibrium between proline synthesis and degradation.

Auxin provides a plausible regulatory link in this equilibrium. Exogenous application of IAA to heat-stressed rice stimulated the ornithine pathway via *OsOAT* upregulation, raising proline levels (Verslues and Sharma, 2010). However, the same hormone drives energy-intensive cell elongation and differentiation, processes that ultimately consume proline through catabolic oxidation. Proline feeds carbon into the tricarboxylic-acid cycle and supplies reducing equivalents to mitochondria, supporting ATP generation and redox balance (Verslues & Sharma, 2010; Sharma et al., 2011). This dual role requires tight spatial and temporal regulation. Therefore, we suggest that exogenous or microbial IAA initially triggers proline biosynthesis to buffer oxidative stress but later promotes proline catabolism as growth resumes. This metabolic shift fuels auxin-driven tissue regeneration and sustains NAD□/NADH homeostasis during rapid cell division. In young, actively growing tissues, proline catabolism may dominate to power thermomorphogenic elongation, while in older tissues, proline accumulation supports osmoprotection and programmed senescence (Pisuttu et al., 2024). Thus, microbial IAA and proline operate as coupled regulators of energy and development under heat stress, a relationship illustrated conceptually (Box 1).

### What mechanisms connect auxin signaling with proline metabolism and energy turnover?

Auxin signaling regulate thermomorphogenic growth through a multilayered network that integrates transcriptional regulation, protein stabilization, redox control, and hormonal crosstalk. Several molecular modules collectively connect auxin perception to proline metabolism and energy turnover, allowing the plant to balance protection and recovery during heat stress.

#### TIR1/AFB-Aux/IAA-ARF cascade stabilized by HSP90

The canonical auxin signaling pathway, mediated by TIR1/AFB co-receptors and Aux/IAA-ARF transcriptional complexes, plays a central role in linking hormonal perception to proline metabolism. Under heat stress, the molecular chaperone HSP90, together with its co-chaperone SGT1, stabilizes the TIR1 receptor complex, preventing its proteasomal degradation (Wang et al., 2016). This stabilization sustains auxin sensitivity even when biosynthesis is transiently reduced, enabling downstream transcriptional control of stress-responsive genes. Among these targets are *P5CS*, which encodes a key enzyme for proline biosynthesis, and *ProDH*, which governs proline catabolism (Verslues and Sharma, 2010; Sharma et al., 2011). Through ARF-dependent regulation of these genes, the TIR1/AFB cascade can toggle metabolic flux between anabolic and catabolic states, coordinating proline turnover with the developmental demands of thermomorphogenesis. Therefore, the HSP90-TIR1 axis not only preserves auxin signal under heat stress but also acts as a molecular conduit linking hormone perception to redox metabolism.

#### MAP kinase and heat shock modules

Heat-induced mitogen-activated protein kinases (MPKs) integrate environmental signals into the auxin network. Under elevated temperatures, MPK3/4/6 phosphorylate the auxin repressor *IAA8*, stabilizing it and transiently repressing ARF activity (Kim et al., 2024). This modulation delays auxin-responsive gene expression until cellular conditions favors growth resumption. In this context, MPK-mediated control of *IAA8* may function as a molecular timer, regulating the transition between proline accumulation and catabolism. During early stress, ARF inhibition promotes *P5CS*-driven proline synthesis for osmoprotection, as MPK activity subsides and IAA8 is degraded, *ProDH* expression increases, supporting energy release and cell elongation. This oscillatory regulation ensures that proline dynamics are synchronized with stress progression, preventing premature energy expenditure and oxidative overload.

#### Mitochondrial redox regulators: SSR1 and HSCA2

Proline oxidation links auxin signaling to mitochondrial energy metabolism. The enzymes ProDH and P5CDH transfer electrons into the mitochondrial electron transport chain (mETC), coupling proline catabolism to ATP generation and redox homeostasis (Verslues and Sharma, 2010). Under heat stress, the integrity of this system is safeguarded by SSR1 and HSCA2, mitochondrial chaperones involved in Fe-S cluster assembly and electron transfer regulation (Han et al., 2021; Zhang et al., 2025). Mutations in these genes disrupt mETC function, leading to proline accumulation, ROS production, and impaired growth recovery (Xia et al., 2025). Auxin signaling may interface with these mitochondrial regulators through redox feedback loops: auxin-induced proline oxidation supplies reducing equivalents that influence TCA cycling, while mitochondrial ROS can modulate auxin transport and sensitivity. Hence, the SSR1-HSCA2 system acts as a redox checkpoint, aligning auxin-driven metabolic demand with the energetic capacity of the cell.

#### Cross-hormonal modulation by ABA and ethylene

Auxin rarely operates in isolation. Hormonal crosstalk with abscisic acid (ABA) and ethylene refines the balance between proline synthesis and degradation. ABA promotes P5CS1 expression and osmoprotective proline accumulation during early stress, whereas ethylene can antagonize or synergize with auxin to modulate growth responses depending on tissue context (Lehr et al., 2022). Although correlations between proline and ABA have been observed, genetic studies reveal partial independence, P5CS1 can still be induced under osmotic stress in ABA-deficient backgrounds (Verslues and Sharma, 2010). This suggests that microbial IAA may complement or substitute ABA signaling during heat stress, maintaining proline biosynthesis when ABA pathways are saturated or repressed. In turn, ethylene and auxin co-regulate cell wall loosening and elongation, linking metabolic energy release from proline catabolism with morphogenetic outcomes (Hare, 2003). Through such cross-hormonal integration, the auxin-proline axis becomes embedded within a broader network that translates environmental cues into coordinated developmental responses.

Together, these interconnected modules TIR1/AFB stabilization, MPK-IAA8 regulation, mitochondrial redox buffering, and hormonal crosstalk compose a dynamic feedback network through which microbial IAA fine-tunes proline metabolism. By modulating the redox and transcriptional landscape of the plant cell, microbial auxin ensures that energy flux, stress protection, and growth recovery are synchronized during thermomorphogenesis. This mechanistic integration provides a foundation for the proposed regulatory window model, wherein microbial IAA acts as a flexible metabolic signal bridging stress defense and developmental plasticity.

### Are there testable hypotheses linking microbial auxin signaling and proline-dependent thermomorphogenesis?

On a mechanistic level, microbial IAA appears capable of rewiring the energetic and redox landscape of the plant cell, yet the precise mechanisms linking microbial auxin input to proline metabolism and thermomorphogenic growth remain unresolved. Integrating the evidence presented above, we propose four complementary hypotheses that can guide future experiments.

#### Hypothesis 1: Auxin perception determines the kinetics of the proline cycle

Auxin perception through the TIR1/AFB-Aux/IAA-ARF module likely governs the timing of the transition from proline accumulation to proline catabolism during heat stress. Under elevated temperature, HSP90 and SGT1 stabilize TIR1, preventing proteasomal degradation and sustaining auxin sensitivity even as biosynthesis declines (Wang et al., 2016). Early in stress, limited ARF activity promotes *P5CS*-driven proline synthesis for osmoprotection (Verslues and Sharma, 2010; Kishor et al., 2022), whereas subsequent Aux/IAA degradation activates *ProDH* expression, providing reducing power for growth recovery (Sharma et al., 2011). Testing this hypothesis could involve comparing *P5CS* and *ProDH* expression dynamics in wild-type and tir1/afb or arf7/19 mutants inoculated with IAA-producing microbes under heat stress to determine whether auxin perception dictates the temporal coordination of these pathways.

#### Hypothesis 2: Mitochondrial redox regulators couple proline oxidation to auxin-driven recovery

A second layer of control may occur at the mitochondria, where the chaperones SSR1 and HSCA2 maintain Fe-S-cluster assembly and electron flow within the respiratory chain (Han et al., 2021; Zhang et al., 2025). Because *ProDH*-mediated proline oxidation provides reducing equivalents to the mETC (Verslues and Sharma, 2010), defects in these proteins elevate ROS and impede thermomorphogenic recovery (Xia et al., 2025). We propose that SSR1 and HSCA2 act as a redox checkpoint linking auxin-induced metabolic demand with the cell’s energetic capacity. This could be examined by quantifying ROS accumulation, proline turnover, and elongation in ssr1 and hsca2 mutants compared with wild type following microbial-IAA exposure.

#### Hypothesis 3: Heat and auxin dose define a regulatory window for sequential *P5CS* and *ProDH* activation

Proline metabolism responds non-linearly to combined heat and hormonal inputs. Low microbial or endogenous IAA levels favour *P5CS* induction and proline accumulation (You et al., 2012; Kishor et al., 2022), whereas sustained or higher auxin inputs accelerate *ProDH* activation, initiating energy turnover (Hare et al., 2003). This shift parallels the progression from protection to recovery during thermomorphogenesis (Casal and Balasubramanian, 2019; Koini et al., 2009). The hypothesis predicts a “regulatory window,” measurable as the interval between *P5CS* and *ProDH* activation across auxin-dose and temperature gradients, which can be monitored using dual *P5CS1::LUC* and *ProDH2::LUC* reporter lines.

#### Hypothesis 4: Microbial IAA acts as an external modulator that retunes the timing of the proline switch

Many thermotolerant microbes, including Bacillus, Pseudomonas, and Trichoderma, sustain IAA production at ≥ 35 °C (Ali et al., 2011; Mukhtar et al., 2020; Singh et al., 2021). We hypothesize that this exogenous auxin supply supplements the plant’s own pool, shortening the protective phase of proline accumulation and advancing growth recovery. IAA-producing strains enhance heat tolerance and biomass accumulation (Ahmad et al., 2023; Dlamini et al., 2025), whereas IAA-deficient mutants confer weaker protection (Ismail et al., 2020). This hypothesis can be tested by comparing proline dynamics and elongation in plants inoculated with producing versus IAA-deficient strains under identical heat regimes to evaluate whether microbial IAA quantitatively retunes host metabolic timing.

## Conclusion

There is increasing evidence that microbial auxins influence plant stress metabolism, yet the mechanisms linking microbial IAA to proline dynamics and thermomorphogenic growth remain unresolved. We posit that microbial IAA acts as a metabolic integrator that connects auxin signalling with redox and energy pathways through coordinated regulation of proline biosynthesis and catabolism. This relationship likely extends beyond canonical transcriptional control, involving post-translational and redox-based feedback via the HSP90-TIR1, MPK-IAA8, and SSR1-HSCA2 modules. The hypotheses outlined here provide a framework to test how microbial auxins create a regulatory window that synchronizes stress protection with growth recovery. Understanding this coordination between microbial IAA production and plant metabolism will open new avenues for exploring microbial contributions to plant thermotolerance and adaptive resilience.

## Acknowledgement

We would like to thank Assia Lachhab for assistance with figure preparation.

## Author contributions

AMK and JW planned and designed the perspective together and both contributed to its writing.

## Data availability

Data sharing is not applicable to this article as no new data were created in this study.

## Key Developments Box

Box 1. Key developments in understanding auxin-proline crosstalk and thermotolerance

1. Kan et al. (2023) uncovered a multilayered temperature-sensing network in plants, revealing how PIF4 and PIF7 are translationally and transcriptionally activated under heat. This study strengthened the mechanistic link between temperature cues, auxin biosynthesis, and early thermomorphogenic responses, establishing a foundation for understanding how metabolic processes including proline turnover are hormonally integrated.
2. Wang et al. (2016) demonstrated that HSP90 stabilizes the auxin co-receptor TIR1 under elevated temperature, uncoupling auxin perception from biosynthesis and ensuring sustained auxin signaling during heat stress.
3. Kim et al. (2024) identified phosphorylation of the auxin repressor IAA8 by heat-responsive MAP kinases (MPK3/4/6), revealing a temperature-dependent timing mechanism that transiently restrains ARF activity during early stress.
4. Han et al. (2021) and Zhang et al. (2025) highlighted the critical roles of the mitochondrial chaperones SSR1 and HSCA2 in maintaining Fe-S cluster assembly and electron transport under stress. These findings position mitochondrial quality control as a central determinant of whether proline oxidation can effectively support thermomorphogenic recovery.

---

**Fig. 1.**
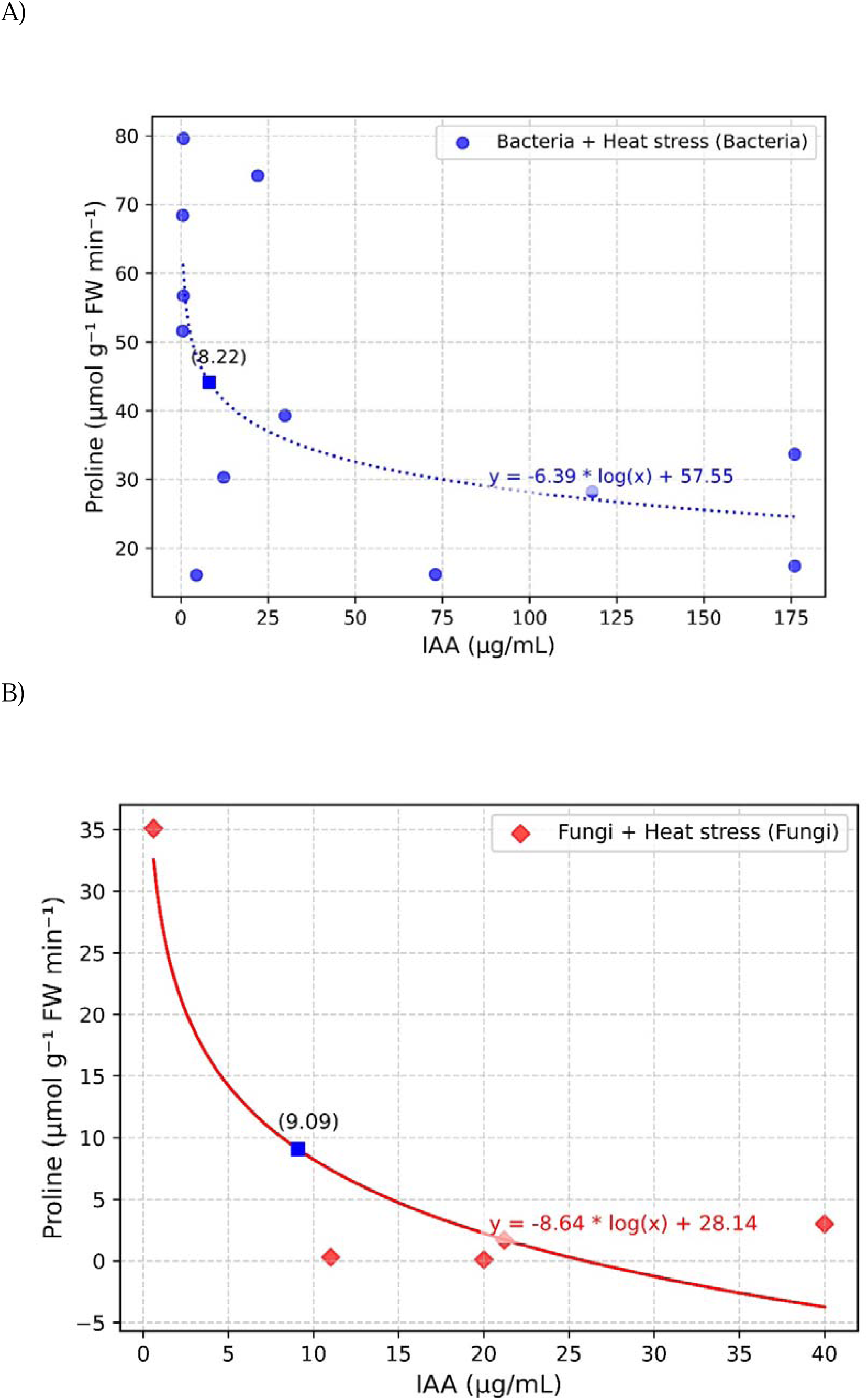
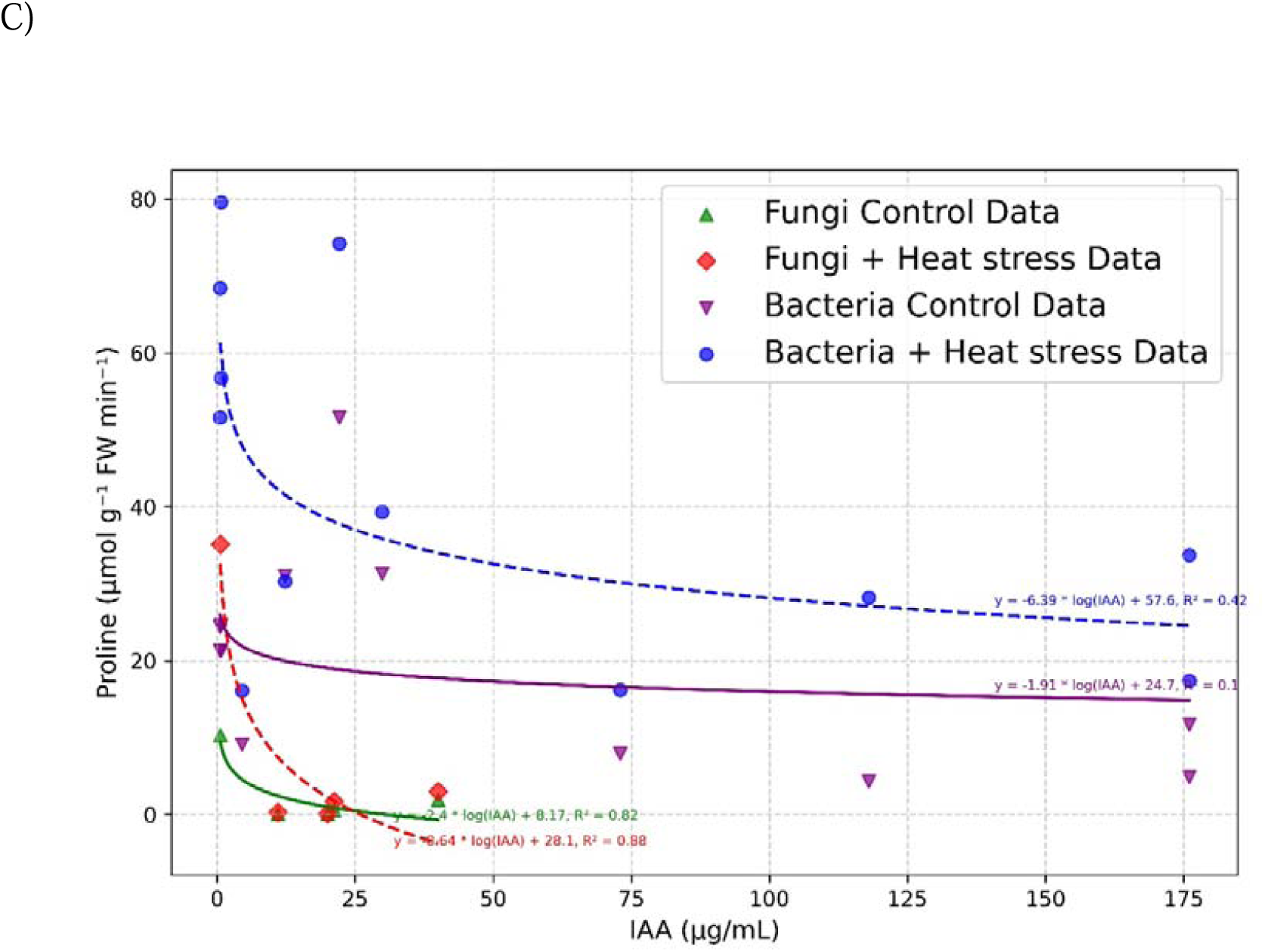
Relationship between exogenous IAA concentration and proline concentration in cool seasonal vegetables and pulses under heat stress. A) The IAA was produced by bacteria {(blue, with heat stress p = 0.03, n = 12) B) fungi (red with heat stress p = 0.2, n = 5). C) Combined plot of bacteria and fungi under heat stress and controlled conditions (purple is bacteria in control condition p = 0.02, n = 12 and green is fungi control condition p = 0.03, n = 5) The best-fit lines were logarithmic regressions (y = m.log(IAA) + b), with test statistics included on the graph with the threshold (blue box) of bacteria and Fungi under heat stress.

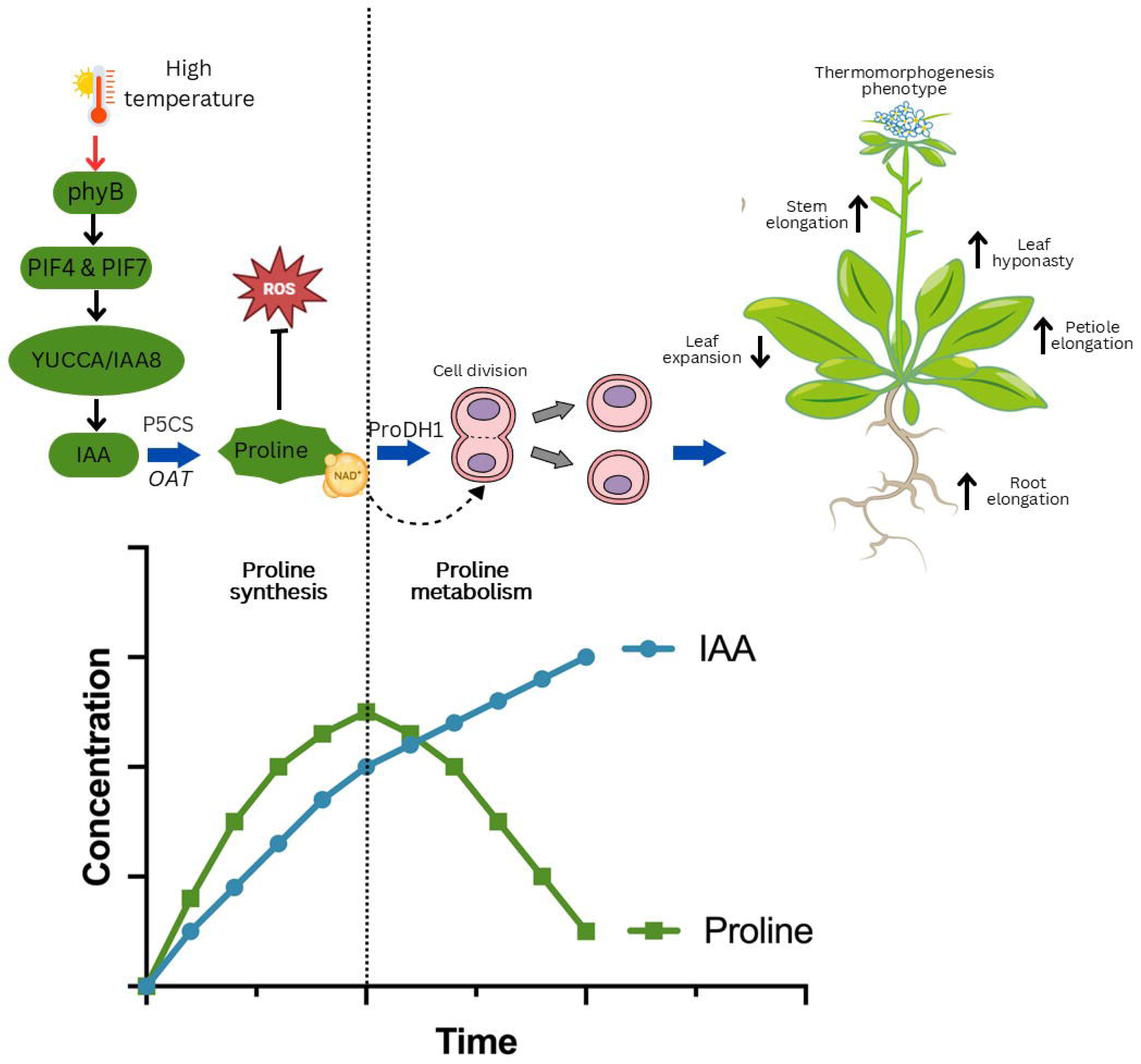

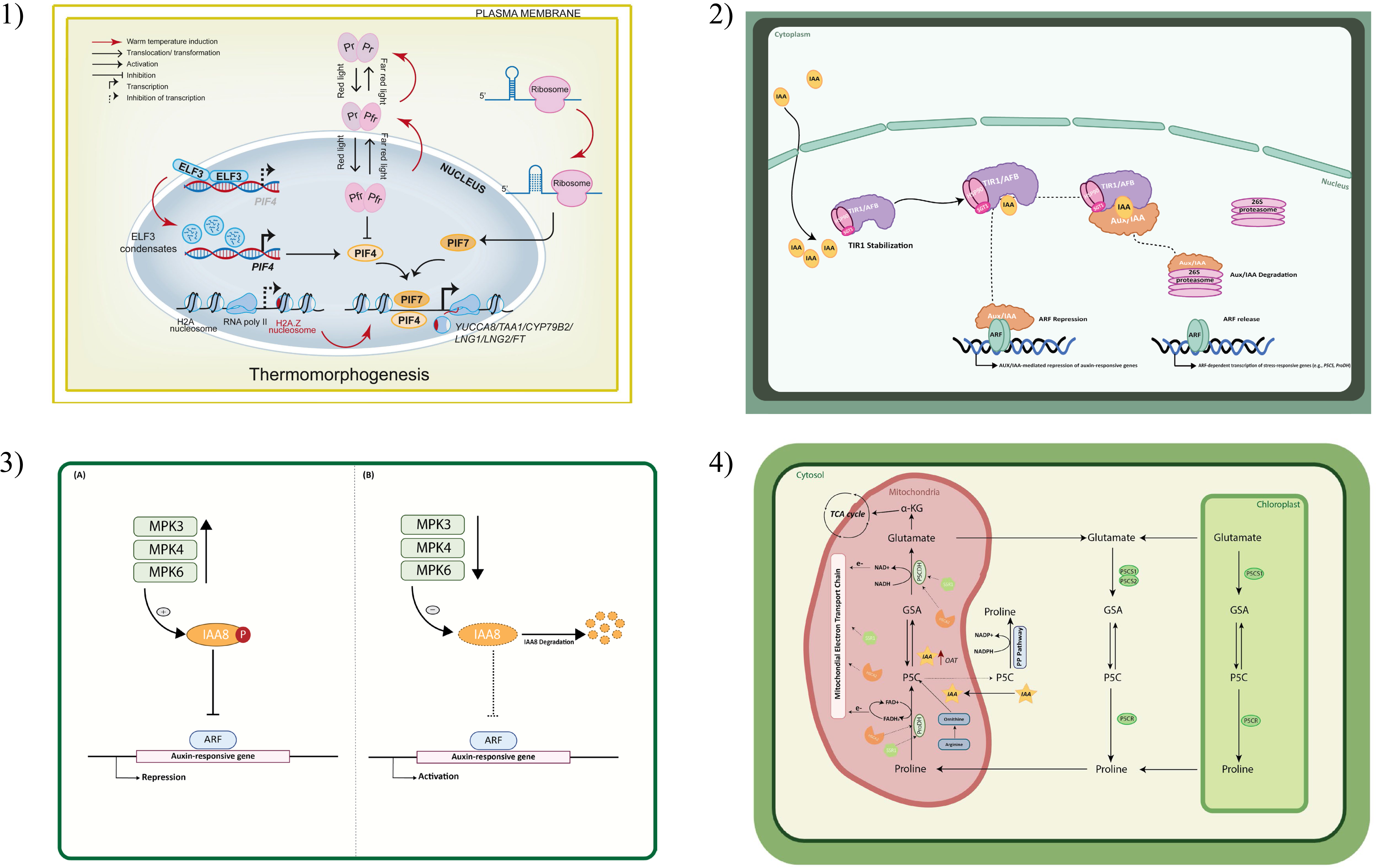

